# KAT7 is a therapeutic vulnerability of *MLL*-rearranged acute myeloid leukemia

**DOI:** 10.1101/2020.04.25.054049

**Authors:** Yan Zi Au, Muxin Gu, Etienne De Braekeleer, Jason Yu, Swee Hoe Ong, Malgorzata Gozdecka, Xi Chen, Konstantinos Tzelepis, Brian J.P Huntly, George Vassiliou, Kosuke Yusa

## Abstract

Histone acetyltransferases (HATs) catalyze the transfer of an acetyl group from acetyl-CoA to lysine residues of histones and play a central role in transcriptional regulation in diverse biological processes. Dysregulation of HAT activity can lead to human diseases including developmental disorders and cancer. Through genome-wide CRISPR-Cas9 screens, we identified several HATs of the MYST family as fitness genes for acute myeloid leukaemia (AML). Here we investigate the essentiality of lysine acetyltransferase KAT7 in AMLs driven by the *MLL-X* gene fusions. We found that KAT7 loss leads to a rapid and complete loss of both H3K14ac and H4K12ac marks, in association with reduced proliferation, increased apoptosis and differentiation of AML cells. Acetyltransferase activity of KAT7 is essential for the proliferation of these cells. Mechanistically, our data propose that acetylated histones provide a platform for the recruitment of MLL-fusion-associated adaptor proteins such as BRD4 and AF4 to gene promoters. Upon KAT7 loss, these factors together with RNA polymerase II rapidly dissociate from several MLL-fusion target genes that are essential for AML cell proliferation, including *MEIS1, PBX3* and *SENP6*. Our findings reveal that KAT7 is a plausible therapeutic target for this poor prognosis AML subtype.

## Introduction

Acute myeloid leukemia (AML) is an aggressive malignancy of haematopoietic stem and progenitor cells. Translocations involving the mixed lineage leukemia gene (*MLL* or *KMT2A*) characterize an aggressive subtype of the disease and are associated with a poor prognosis ^1–3^. Currently, chemotherapy based on a combination of anthracyclines and purine analogues is the standard of care for AML, yet patients with leukemias driven by *MLL-X* gene fusions commonly become refractory to such treatments ^4^. Advances in our understanding of the molecular pathogenesis of acute leukemias driven by *MLL-X* fusion oncogenes have led to the development of new drugs, including inhibitors of bromodomain proteins ^5–7^, DOT1L ^8,9^ and MENIN ^10^; however, none of these have demonstrated a significant clinical benefit for patients as yet. In order to identify new therapeutic approaches in AML, we and others have performed genome-wide CRISPR-Cas9 fitness screens in different subtypes of the disease including the *MLL-X* subtype^11,12^. Through our own screen data, we recently showed that GCN5 (KAT2A) and SRPK1 are novel vulnerability in AML and that sensitivity to SRPK1 inhibition is particularly associated with AML driven by *MLL-X* fusion oncogenes ^11,13^. Systematic detailed analysis of our CRISPR screen dataset revealed that several members of the MYST family of histone acetyltransferases (HATs) are required for the survival of AML cell lines, with KAT7 displaying strong essentiality for AMLs driven by *MLL-X* oncogenes.

HATs are classified based on structural and sequence homology into distinct groups including the p300/CBP, MYST and GCN5 families ^14^. They play a central role in transcriptional regulation through their function to catalyze the transfer of acetyl groups from acetyl-CoA to the ε-amino group of lysine residues of histones. Dysregulation of HATs is known to be associated with human diseases including developmental disorders and cancer ^14–16^. The MYST family of HATs, characterized by their conserved MYST catalytic domain, includes KAT5 (TIP60), KAT6A (MOZ and MYST3), KAT6B (MORF and MYST4), KAT7 (HBO1 and MYST2) and KAT8 (MOF)^15,16^. While *KAT6A* and *KAT6B* are known targets of chromosomal translocations that drive AML^17^–^21^ and KAT8 can play a role in some cases of AML ^22^, it is currently not known whether KAT7 plays a role in leukemogenesis. As KAT7 lacks chromatin binding domains, it relies on forming complexes with a scaffold protein (JADE or BRPF), along with EAF6 and ING4/5, for efficient histone acetylation ^23^. The choice of the scaffold protein primarily determines target histones, namely H3 or H4. The KAT7-BRPF acetylates histone H3 tails (K14 and K23)^23^–^26^, whereas KAT7-JADE complexes acetylates histone H4 tails (K5, K8, and K12)^27^, respectively. In development, abolishing KAT7 results in major reduction of H3K14ac in both mouse fetal liver erythroblasts ^24^ and mouse embryo ^25^; the latter associates with lethality.

Here we investigate the molecular mechanism underlying the requirement of KAT7 in *MLL-X*-driven AMLs. We report that loss of KAT7 leads to reduced proliferation and enhanced apoptosis and differentiation. We also show that KAT7 is essential for maintaining the transcriptional program driven by MLL-AF9, through the recruitment of BRD4 and other MLL-fusion associated proteins such as AFF1 to the promoters of a subset of MLL-AF9 target genes. Together, our findings propose KAT7 as a plausible therapeutic target in *MLL-X* AML and also provide novel mechanistic insights into the molecular basis of MLL-X-driven transcriptional dysregulation.

## Materials and Methods

### Cell culture

MOLM-13, MV4-11, THP-1, HL-60 and K562 were cultured in RPMI 1640 medium (Thermo Fisher) supplemented with 10 % FBS (Thermo Fisher). Nomo-1 was cultured in RPMI 1640 supplemented with 10 % FBS, 1 % penicillin/streptomycin (Thermo Fisher), 1 % sodium pyruvate (Thermo Fisher), 1 % glucose (Sigma). OCI-AML2 and OCI-AML3 were cultured in MEM-α (Lonza) with 20 % FBS and 1 % GlutaMax (Thermo Fisher). All these cell lines stably express Cas9 (ref. 11). To maintain Cas9 expression, blasticidin (Thermo Fisher) was added to MOLM-13, MV4-11, THP-1, Nomo-1, HL-60, OCI-AML2 and OCI-AML3 culture at 10 μg/ml, and K562 at 15 μg/ml. 293FT cells (Thermo Fisher) were cultured in DMEM (Thermo Fisher) supplemented with 10 % FBS and 1 % GlutaMax. All cell lines were incubated at 37 °C with 5 % CO_2_ and confirmed to be mycoplasma negative.

### Plasmid construction

#### gRNA design and cloning

gRNAs from the genome-wide CRISPR-library used in the initial screens ^11^ were selected for validation. Additional gRNAs were designed using the Wellcome Sanger Institute’s Genome Editing (WGE) (http://www.sanger.ac.uk/htgt/wge/)^28^. Only gRNAs that targeted exons present in all putative transcripts and had no off-target hits with less than 3-nucleotide mismatch to their on-target sequence were selected. All gRNAs were cloned into the Bbsl site of pKLV2-U6gRNA5(BbsI)-PGKpuro2ABFP-W ^11^. gRNAs used in this study are listed in Table S2.

#### gRNA-resistant wild-type and E508Q mutant KAT7 expression vector

A T2A-fused wild-type (WT) KAT7 cDNA fragment resistant to both gKAT7 (5) and gKAT7 (A10) was purchased from IDT and cloned into the BsrGI site of pKLV-EF1aGFP using a NEBuilder HiFi kit (NEB), resulting in pKLV-EF1aGFP2AKAT7-W. To introduce E508Q substitution, the region between the SrfI and BsaBI sites of the KAT7 ORF was replaced with a gBlock fragment containing the substitution using NEBuilder HiFi kit, resulting in pKVL-EF1aGFP2AKAT7E508Q-W. Sequences were verified by Sanger sequencing.

#### KAT7 degron-AID system

The region between the BamHI and NotI of pKLV-EF1aGFPKAT7-W was replaced with a gBlock fragment appended with the degron tag using NEBuilder HiFi kit, resulting in pKLV-EF1aGFP2AKAT7-linkermAID-W. A T2A-fused *Oryza sativa* TIR1 (OsTIR1) fragment was purchased as a gBlock fragment and cloned into the BsrGI site of pKLV-EF1amCherry, resulting in pKVL-EF1a-mCherry-2A-OsTIR1-W. Sequences were verified by Sanger sequencing.

### Lentivirus production and transduction

Lentivirus was produced as described previously (31). AML and CML cell lines were transduced by adding lentivirus and 8 μg/ml polybrene (Millipore) to 3.0 x 10^4^ cells per well of a 96-well plate or 1 x 10^6^ cells per well of a 6-well plate and incubated at 37°C for 22 h. Viral supernatant was then replaced with fresh culture media and the cells were passaged for further culture.

### Cell sorting and flow cytometry analyses

Cell sorting was performed using Mo-Flo XDP or BD INFLUX under containment-level 2 conditions. FACS analyses were performed using BD LSRFortessa and raw files were analysed using FlowJo.

### Proliferation

Cells were first transduced with gRNA-expressing lentivirus in 96-well plates as described above. The percentage of BFP-positive (knock-out) cells was determined for each well every 2 days between day 4 and day 14 post transduction using flow cytometry. Culture media were refreshed every 2 days. Prior to FACS analysis, cells were fixed in 4 % paraformaldehyde in phosphate-buffered saline (PBS) for 10 minutes and re-suspended in 1 % bovine serum albumin in PBS. Values were normalized to the BFP-positive percentage on day 4.

### Differentiation and apoptosis assays

Cells were transduced with gRNA-expressing lentivirus in 6-well plates as described above. BFP-positive (knock-out) cells were collected by cell sorting 3 days post transduction and further cultured until analysis. For differentiation analysis, 2.3 x 10^5^ cells were harvested 7 days post transduction, washed in PBS and staining buffer (2 % FBS in PBS), and stained with APC-conjugated CD11b (1.15 μl antibody per 100 μl staining buffer per test, 17-0118, eBioscience) or APC-conjugated mouse IgG1κ isotype control (17-4714, eBioscience) on ice for 30min in the dark. The cells were then washed twice with staining buffer and analysed by FACS. For apoptosis assays, 2.3 x 10^5^ cells were analysed 9 days post transduction using Annexin V Apoptosis Detection Kit APC (88-8007, eBioscience) according to the manufacturer’s instruction.

### Generation of Auxin Inducible Degron (AID) KAT7 protein degradation system and treatment with indole-3-acetic acid (IAA)

MOLM-13 cells were first transduced with lentivirus produced with pKLV-EF1aGFP2AKAT7_linkermAID-W and GFP-positive cells were sorted. The cells were then transduced with lentivirus produced with pKLV2-U6gKAT7(A10)-PGKpuro2ABFP-W to knock out the endogenous *KAT7* gene and sorted for BFP-positive cells to enrich for the knock-out (KO) population. GFP-BFP-double positive cells were further transduced with lentivirus produced with pKLV-EF1a-mCherry-2A-OsTIR1-W and sorted for mCherry-positive cells, resulting in the generation of MOLM-13 KAT7-AID cells. Indole-3-acetic acid (IAA; Sigma) was dissolved in water at a concentration of 500 mM. MOLM-13 KAT7-AID cells were treated with 500 μM IAA for 24 h for ChIP-qPCR experiments and for 48 h for differentiation assays.

### Western blot analysis

Cells were harvested by centrifugation and washed twice in PBS. Cell pellets were then re-suspended in NuPAGE LDS sample buffer (ThermoFisher), NuPAGE sample reducing agent (ThermoFisher) at a concentration of 1 x 10^6^ cells/100 μL. The lysates were heated at 95 °C for 5 min and vortexed at room temperature for 10 min. Equal volumes of the lysates were loaded into each well of Bis-Tris precast polyacrylamide gels (Thermo Fisher). Antibodies used are listed in Table S3.

### In vivo mouse work

MOLM-13 cells expressing Cas9 and luciferase ^11^ were transduced with lentivirus for gRNA-mediated knock-out of KAT7 using gKAT7 (A10) or empty gRNA control. BFP-positive cells were sorted 2 days post transduction and cultured. 5 x 10^5^ cells were injected via the tail vein of immunocompromised NSGW41 male mice (derived by breeding NSG and KITW41 animals to homozygosity, i.e. *Kit^W41/W41^, Prkdc^-/-^*, and *Il2rg^-/-^* or *Il2rg^-/Y^*) 3 days post transduction. Five mice were used in each treatment group. Bioluminescence imaging was performed by injection of 200 μL (2 mg/mL) D-luciferin (BioVision) per mouse. Quantification of bioluminescence was performed using the IVIS Spectrum In Vivo Imaging System (Perkin-Elmer), with Living Image version 4.3.1 software (PerkinElmer) according to the manufacturer’s instructions. Log-rank (Mantel-Cox) test was used to compare survival between mouse groups. All animal studies were carried out at the Wellcome Sanger Institute under UK Home Office License PBF095404.

### ChIP-seq and ChIP-qPCR

Cells were crosslinked with 1 % Formaldehyde (ThermoFisher) at room temperature for 5 min (histones ChIP) or 10 min (non-histone ChIP), and quenched with 0.125M Glycine (Sigma) at room temperature for 5 min. Cross-linked cells were washed twice with ice-cold PBS before sequential lysis with LB1 (50 mM Hepes, 140 mM NaCl, 1 mM EDTA, 10 % glycerol, 0.5 % NP-40, 0.25 % Triton X-100), LB2 (10 mM Tris-HCl, 200 mM NaCl, 1 mM EDTA, 0.5 mM EGTA) and LB3 (10 mM Tris-HCl, 100 mM NaCl, 1 mM EDTA, 0.5 mM EGTA, 0.1 % Na-Deoxycholate, 0.5 % N-lauroylsarcosine) on ice for 10 min each. Lysed samples were then sonicated for 10 and 11 cycles for MOLM-13 and OCI-AML3 respectively, using the Bioruptor Pico instrument (Diagenode). Triton-X was added to the sonicated samples to a final concentration of 1% before centrifugation at 20,000 x g at 4 °C for 10 min. 10 % (by volume) of each sample was kept as input. Lysates were incubated with antibody (Table S2) at 4 °C overnight with rotation. Dynabeads Protein A/G (ThermoFisher) were added the next day and incubated at 4 °C for 4 h with rotation. ChIP samples were washed with RIPA wash buffer (50 mM Hepes, 500 mM LiCl, 1 mM EDTA, 1 % NP-40, 0.7 % Na-Deoxycholate) 3 times, followed by wash with annealing buffer (TE, 50 mM NaCl). Elution buffer (1 % SDS, 50 mM Tris-HCl, 10 mM EDTA) was added to each ChIP sample and the samples were then heated at 65 °C for 30 min. Beads were subsequently removed and samples heated at 65 °C overnight to reverse cross-linking whilst supplemented with 0.2 mg/mL RNase A (ThermoFisher). 0.2 mg/mL Proteinase K (Life Technologies) was added the next day to digest proteins by incubating and shaking at 450 rpm at 65 °C for 4 h. ChIP-DNA was purified by PCR purification Kit (Qiagen). PBS and LB1, LB2, LB3 and RIPA wash buffers were all supplemented with Sodium Butyrate (Sigma), cOmplete EDTA-free protease inhibitor cocktail (Roche) and PMSF (Sigma) immediately before use. Sequencing was performed on Illumina HiSeq v4 platform using 75-bp paired-end sequencing.

For qPCR, KAPA SYBR Fast qPCR kit was used following the manufacturer’s protocol. % Input was calculated as 100 x 2^Δct^, where ΔCt = ct_Normalized input_− ct_Normalized input_ = Ct_Input_ log_2_ (input dilution factor); input dilution factor = fraction of input saved relative to each IP x dilution of input for qPCR. Sequences of primers used in ChIP-qPCR are listed in Table S4.

### ChIP-seq data processing and analysis

Reads from ChIP-seq samples and input were mapped to the human genome GRCh38 using Burrows-Wheeler Aligner (BWA) version 0.7.17 (ref. 29) on default parameters. Duplicates were marked by Samtools mkdup ^30^. Peaks (broad and narrow) were called by MACS2 version 1.4.1 (ref. 31) using input DNA as control with parameters --broad-cutoff 0.01 for broad peaks, and −q 0.01 for narrow peaks. Broad and narrow peaks were merged into a union set. Broad peaks that overlap with one or more narrow peaks were removed. Locations of peaks (promoter, exonic, intronic or intergenic) were computed by customized scripts using Ensembl transcript annotation of GRCh38 version 91. Peaks were considered to be associated with promoter(s) if more than a half of the peak region was located within ±2kb from the transcription start site. Promoter occupancies of a transcript were quantified as the highest MACS2 signal among all the peaks within the 2kb window. If multiple isoforms existed, genic promoter occupancies were calculated as the highest signal among isoforms. Public data used was GSM1845161 and GSE83671.

### RNA extraction and RNA-seq processing

RNA was extracted from AML cells with RNeasy Plus Mini Kit (Qiagen) according to manufacturer’s instructions. Sequencing was performed on Illumina HiSeq v4 platform with 75-bp paired-end sequencing.

RNA-seq reads were mapped to the human genome assembly GRCh38 using STAR version 6.43 (ref. 32), tolerating mismatch rate of 0.01 and allowing maximal intron lengths of 50kb. Read counts were calculated by STAR using Ensembl annotation of GRCh38 version 91 and normalized as fragments per kilobase of gene length per million uniquely mapped reads (FPKM). Differential expression analysis was done by DESeq2 (ref. 33) using paired sample design and significant genes were identified using adjusted *p-values* of 0.05 as a cut-off.

RNA-seq and ChIP-seq data are available from Gene Expression Omnibus under accession number GSE133516 (Security Token, svobogacrrchtkd).

### Statistical analysis

Student’s *t*-test was used for statistical testing unless stated otherwise. Mean was calculated from at least three replicates, as indicated in each figure, and error bars represent the standard deviation. *P* values ≤0.05 were considered statistically significant.

## Results

### Loss of *KAT7* induces myeloid differentiation and apoptosis in AML cell lines with MLL fusion oncoprotein

We previously catalogued fitness genes in 5 AML cell lines (MOLM-13, MV4-11, OCI-AML2, OCI-AML3 and HL-60) by genome-scale CRISPR-KO screening and characterized promising therapeutic targets including the histone lysine acetyltransferase KAT2A ^11^. We further looked into genes encoding histone modifying enzymes and found that several histone acetyl transferases (HATs) were required for AML cell proliferation (Fig. 1A). Notably, two of the MYST family HATs, KAT6A and KAT7, were essential for AML cell lines driven by *MLL-X* fusion genes: MOLM-13 (driven by MLL-AF9), MV4-11 (MLL-AF4) and OCI-AML2 (MLL-AF6). In order to validate these fitness defects, we first investigated the effects on proliferation in MOLM-13 and MV4-11 by gRNA-mediated KO and found that *KAT7* loss exhibited a consistent strong anti-proliferative effect in both cell lines (Fig. S1A). We then tested the impact of genetic disruption of *KAT7* on proliferation of all cell lines used in our CRISPR-KO screens and 3 additional cell lines, and found that, consistent with MOLM-13 and MV4-11, *KAT7* loss decreased the proliferation of other *MLL-X* cell lines, namely OCI-AML2, THP-1 (MLL-AF9) and Nomo-1 (MLL-AF9), but not of *MLL*-WT cell lines such as OCI-AML3, HL-60 and K562 (Figs. 1B and S1BC). The proliferation phenotype was associated with increased expression of the myeloid differentiation marker CD11b in MOLM-13 and OCI-AML2 (Fig. 1C). *KAT7* loss also led to an increased apoptosis in *MLL-X* cell lines (Fig. 1D). Collectively, these results indicate that *KAT7* is required for *in vitro* survival and proliferation of leukemic cells driven by *MLL-X* fusion oncogenes. To assess if *KAT7* inactivation has also anti-proliferative effects *in vivo*, we injected MOLM-13 cells previously transduced with a *KAT7* gRNA (*KAT7*-KO) into NSGW41 male mice. Mice transplanted with *KAT7-KO* MOLM-13 cells showed significantly slower AML progression and increased survival, compared to those injected with MOLM-13 expressing a control gRNA (Fig. 1E, F), indicating the requirement of KAT7 for *in vivo* proliferation.

**Figure 1.**
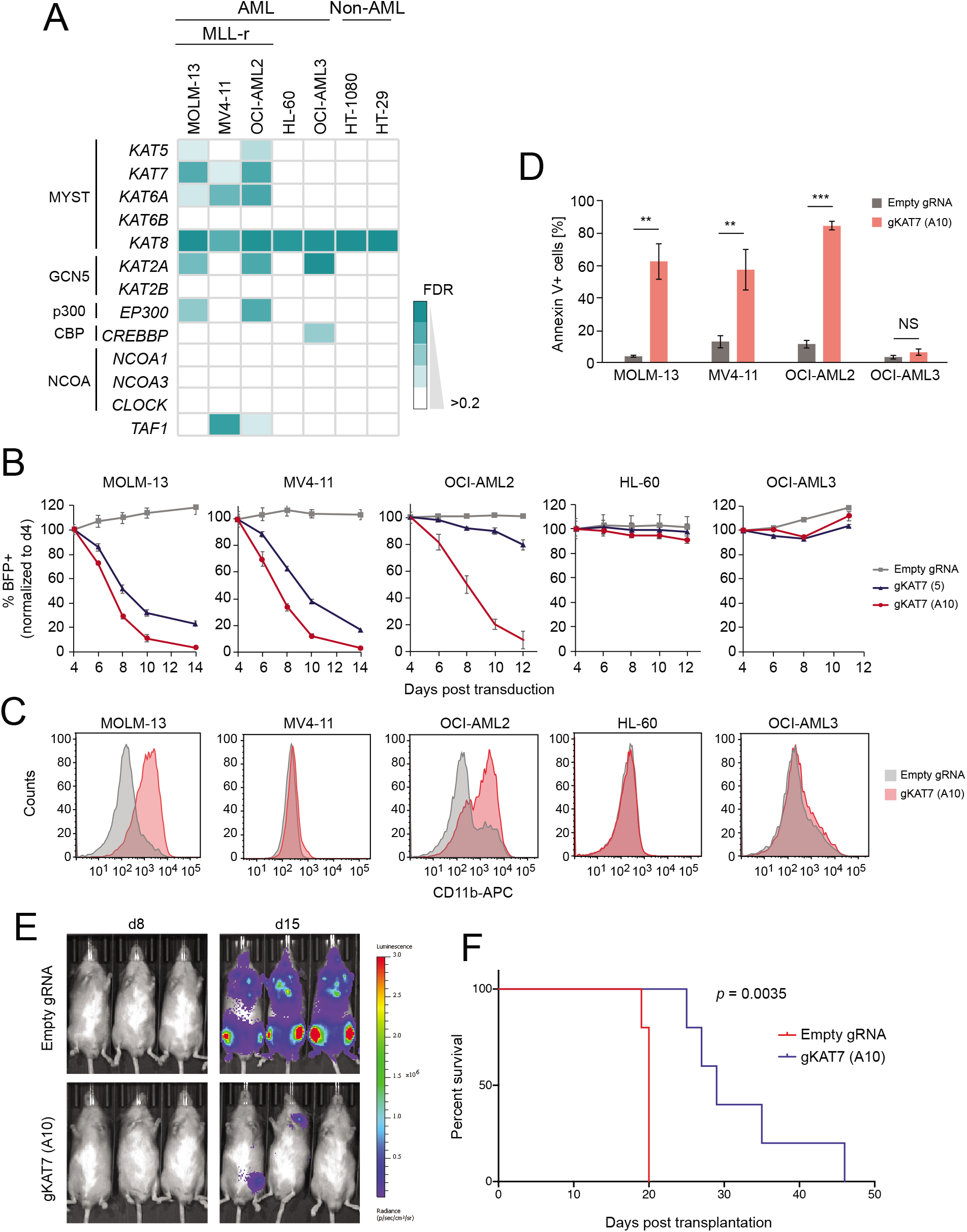
Loss of KAT7 exhibits anti-leukemic effects *in vitro* and *in vivo*. **A)** Fitness effect of HATs from genome-scale CRISPR-KO screening in AML and non-AML cell lines ^11^ based on false discovery rate (FDR) values. **B)** Proliferation of *KAT7* KO using two sgRNAs or empty control in AML cell lines. Percentages of BFP-positive (*KAT7*-KO) cells were assayed at the indicated time point and were normalized to day 4. Data are shown as mean ± SD (n= 3). **C)** CD11b staining in WT (empty gRNA) and *KAT7* KO (gKAT7-A10) cells 7 days post transduction. **D)** Annexin V/PI staining of WT (empty) and *KAT7* KO (gKAT7-A10) cells 9 days post transduction. Data are shown as mean ± SD (n= 3). Two-tailed *t*-test was performed (N.S., not significant; **, *P* < 0.01; ***, *P* < 0.001). **E-F)** Xenograft analysis of *KAT7-KO* MOLM-13. Bioluminescent imaging (**E**) and Kaplan-Meier survival analysis (**F**) were performed (n=5 in each arm).

### KAT7 regulates global H3K14 and H4K12 acetylation

KAT7 is known to form two distinct HAT complexes that acetylate either H3 or H4^23^. We therefore investigated the acetylation status of H3 and H4 tails in *KAT7* KO MOLM-13 cells. Among the potential KAT7 acetylation sites, acetylation of Lysine-14 residue of H3 (K3K14ac) and Lysine-12 of H4 (H4K12ac) were completely abolished upon *KAT7* KO, whereas other acetylation sites were not affected (Fig. 2A). We additionally investigated major histone marks for active transcription, namely H3K4me3, H3K9ac and H3K27ac, and found that *KAT7* loss did not affect the global level of these marks (Fig. 2B). Interestingly, global loss of H3K14ac and H4K12ac upon KAT7 loss was observed in all AML cell lines tested, irrespective of the MLL translocation status or proliferation phenotype (Fig. 2C).

**Figure 2.**
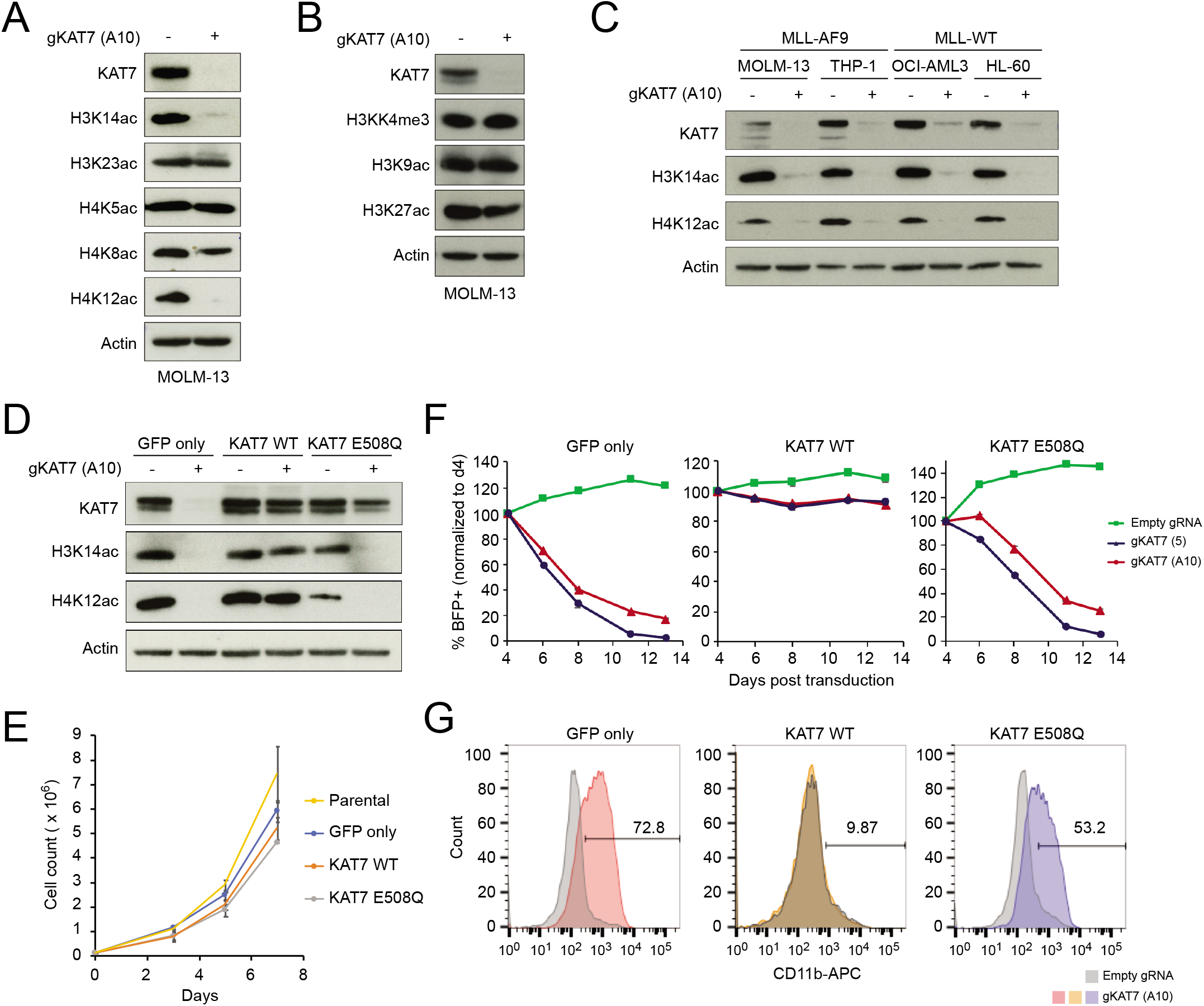
Catalytic activity of KAT7 is required for leukemic maintenance. **A-B)** Western blot analyses of potential KAT7 acetylation sites on H3 and H4 (**A**) and major histone marks associated with active transcription (**B**) in *KAT7-KO* cells. **C)** Western blot analysis of KAT7-mediated acetylation on H3K14 and H4K12 in an additional MLL-AF9+ve cell line, THP-1, and *MLL*-WT OCI-AML3 and HL-60. **D)** Western blot analysis of MOLM-13 expressing exogenous WT-KAT7 or HAT-dead KAT7 (E508Q) with or without endogenous *KAT7* KO by lentiviral gRNA expression. **E)** Proliferation of MOLM-13 cells expressing exogenous KAT7 WT or E508Q mutant as well as endogenous KAT7. **F-G)** Proliferation (**F**) and CD11b staining (**G**) of MOLM-13 expressing exogenous KAT7 WT or E508Q mutant with endogenous KAT7 disrupted by gKAT7.

### Acetyltransferase activity of KAT7 is essential for leukemic maintenance

Since both *MLL-X* and *MLL-WT* AML cell lines showed globally reduced H3K14 and H4K12 acetylation upon *KAT7 KO* but only *MLL-X* AML cell lines showed reduced proliferation, we considered that non-catalytic roles of KAT7 complexes may have played a role in the observed proliferative defect. To rule out such possibility, we performed cDNA reconstitution experiments using gRNA-resistant WT and HAT-dead *KAT7* cDNAs. For the HAT-dead KAT7 mutant, we utilized a previously characterized E508Q MYST domain mutant ^34^. To replace endogenous KAT7 with a cDNA-derived protein, we first expressed a gRNA-resistant *KAT7* cDNA and then knocked out the endogenous *KAT7* by lentiviral transduction of *KAT7* gRNA. Western blot analyses showed that control (GFP only) MOLM-13 cells lost KAT7 as well as acetylation marks on both H3K14 and H4K12 upon the endogenous *KAT7* KO, whereas cells carrying the WT *KAT7* cDNA retained KAT7 protein and both acetylation marks, even after gRNA-targeting of the endogenous *KAT7* gene (Fig. 2D), indicating the appropriate reconstitution of *KAT7* activity. In contrast, cells expressing HAT-dead E508Q mutant KAT7 protein completely lost both acetylation marks after endogenous *KAT7* KO (Fig. 2D). We noted that cells expressing both endogenous KAT7 and exogenous HAT-dead KAT7 showed considerably reduced H4K12ac, although H3K14ac was maintained at the level similar to that of the control cells. This suggest a dominant negative effect of HAT-dead KAT7 protein; however, proliferation of MOLM-13 cells expressing either exogenous KAT7 was not significantly different (Fig. 2E). We then analyzed proliferation of KAT7-reconstituted cells and found that *KAT7*-WT-expressing cells did not show a proliferation defect, ruling out the possibility that off-target effects of *KAT7* gRNAs might have been responsible for the observed phenotype (Fig. 2F, middle panel). In sharp contrast, the cells expressing HAT-dead KAT7 showed proliferation defect(Fig. 2F, right panel), similar to that in endogenous *KAT7* KO cells. This proliferation defect was associated with increased CD11b expression (Fig. 2G). Thus, the cells expressing HAT-dead KAT7 phenocopied the endogenous *KAT7* KO cells in histone acetylation, proliferation and differentiation. These results strongly indicate that KAT7-mediated acetylation plays crucial roles in the maintenance of the leukemic program driven by the MLL-X fusion protein.

### KAT7 loss leads to down-regulation of a specific set of MLL-AF9 target genes

In order to analyze transcriptomic changes upon *KAT7* KO, we performed RNA-seq analyses of cells transduced with lentivirus expressing *KAT7* or control gRNA. We chose MOLM-13 and OCI-AML3 as the *MLL-X* positive and negative cell lines, respectively, in this analysis. On day 3 after transduction, we observed a total of 544 differentially expressed (DE) genes (317 up- and 227 down-regulation) in MOLM-13 (Fig. 3A). The number of DE genes increased further on day 5 to 4093 (1954 up- and 2139 down-regulation). In contrast, the KAT7-independent OCI-AML3 showed only 11 genes on day 3 and 501 on day 5. Consistent with myeloid differentiation observed by flow cytometry (CD11b upregulation, Fig. 1C), gene set enrichment analysis (GSEA) revealed that a gene set consisting of genes upregulated upon myeloid differentiation was significantly enriched on days 3 and 5 (Fig. 3B, left). Concomitantly, the MLL-AF9 target gene set was significantly depleted on day 5 (Fig. 3B, center bottom). Interestingly, although negative enrichment of the MLL-AF9 target gene set was not yet detected on day 3, enrichment of the myeloid differentiation-associated gene set was already evident (Fig. 3B, top), indicating the rapid transcriptomic alteration upon KAT7 loss and thus key roles of KAT7 in maintaining MLL-AF9-driven leukemic program.

**Figure 3.**
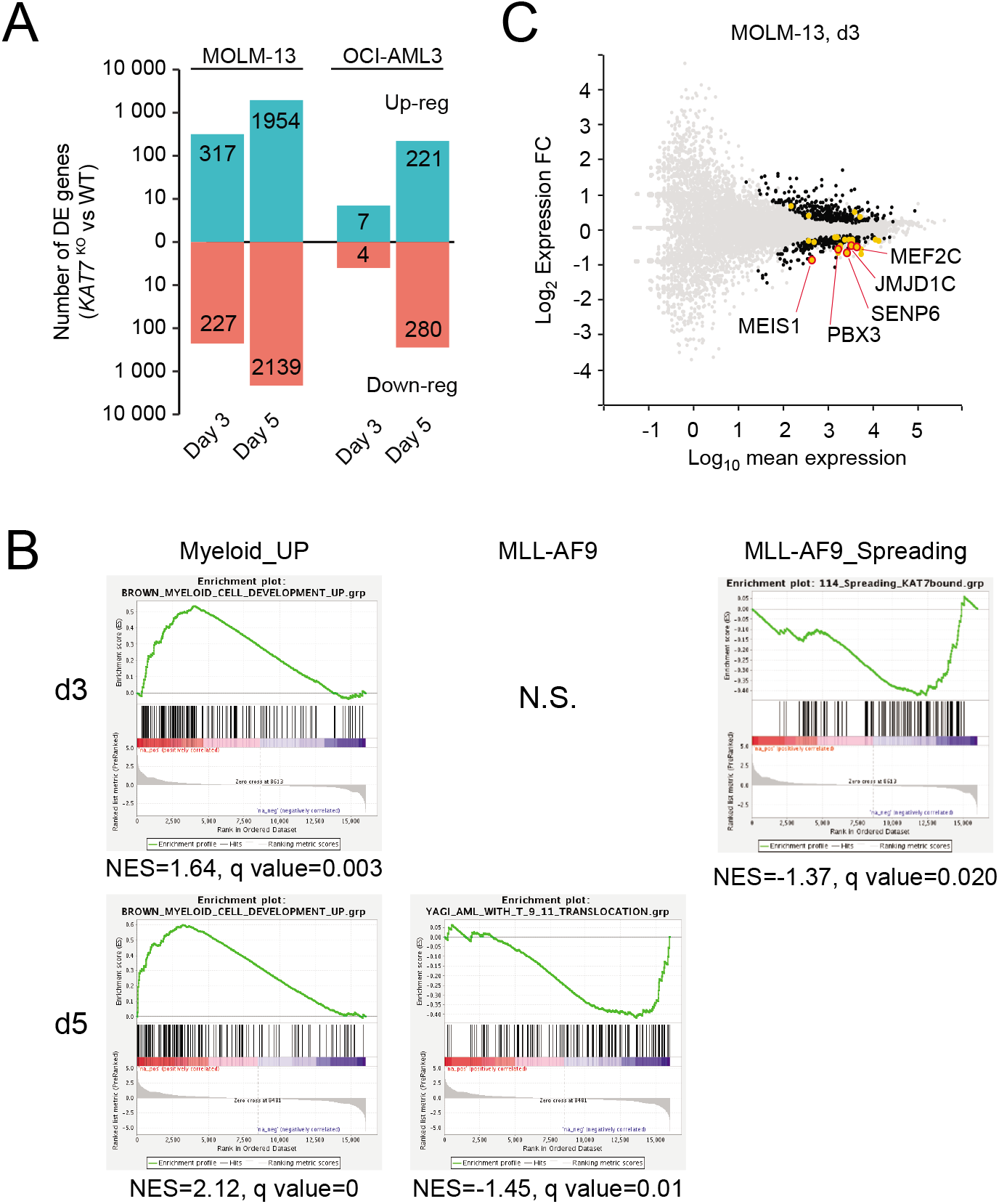
Transcriptomic profiling of *KAT7* KO in MOLM-13 (MLL-AF9) and OCI-AML3 (*MLL* WT) **A)** Numbers of differentially expressed (DE) genes in MOLM-13 and OCI-AML3 on day 3 and day 5 after gRNA-mediated *KAT7* KO. **B)** GSEA on day 3 (top) and day 5 (bottom) transcriptomes in MOLM-13. Gene sets consisting of genes upregulated upon myeloid differentiation (left), MLL-AF9 target genes (centre), or MLL-AF9 spreading genes (right; Table S1) were used. **C)** MA plot of day 3 MOLM-13 comparing *KAT7* WT and KO. Black, DE genes (adj.P<0.05); yellow, MLL-AF9 spreading genes. Key MLL-AF9 spreading genes, namely *MEIS1, PBX3, JMJD1C, SENP6* and *MEF2C* were highlighted.

Through inspection of differentially expressed genes in MOLM-13 on day 3, we identified that a small fraction of the MLL-AF9 target genes, most notably *MEIS1, PBX3, JMJD1C, SENP6* and *MEF2C*, were significantly down-regulated (Fig. 3C). Together with HOXA9, MEIS1 and PBX3 are known to form a complex and promote transcription of HOXA target genes ^35^. JMJD1C and MEF2C are also known to play an important role in MLL-AF9-induced leukemogenesis ^36,37^. In addition, these genes have recently been shown to be among genes, of which MLL-AF9 binding is not limited to their promoter region but spreads into their gene body ^38^. These MLL-X oncoprotein “spreading” genes are collectively more susceptible to DOT1L inhibition than MLL-X “non-spreading” genes ^38^, indicating that spreading genes comprise the core downstream network of the *MLL-X*-driven leukemic program. We further performed GSEA on the day 3 expression profile with a gene set consisting of protein-coding KAT7-bound MLL-AF9 spreading genes (see below, Table S1). Although enrichment of the comprehensive MLL-AF9 target gene set was not detected on day 3 (Fig. 3B, center), this smaller subset was significantly depleted (Fig. 3B, right). Enrichment was not observed in the *MLL*-WT, KAT7-independent OCI-AML3 cell line. These results indicate that KAT7 loss disrupts a leukemia maintenance program through down-regulation of the MLL-AF9 spreading genes, rather than the entirety of MLL-AF9 target genes.

### KAT7 binds to promoters of active genes, especially a subset of MLL-AF9 spreading genes

We next investigated the genomic location of KAT7 binding sites using chromatin-immunoprecipitation followed by sequencing (ChIP-seq) analysis. We again chose MOLM-13 and OCI-AML3 for this analysis and identified 9214 and 60153 KAT7 peaks, respectively (Fig. 4A). KAT7 was most enriched at promoters in both cell lines; 63.3 % of the KAT7 signals in MOLM-13 and 48.5 % in OCI-AML3 were found in promoter regions (±2 kb from TSSs) (Fig. 4A, B, D). Consistent with previously reported ChIP-seq studies ^16,23^,^24^, KAT7 occupancy in the AML cell lines is significantly correlated with the expression levels of target genes (Fig. 4B-E). We then focused our analysis on the MLL-AF9 spreading gene set. These genes are expressed not only in the MLL-AF9+ve MOLM-13 but also in MLL-AF9-ve OCI-AML3 with a significant correlation of their expression levels between the two cell lines (Fig. 4F, r^2^—0.80). As KAT7 marks expressed genes, the majority of the MLL-AF9 spreading genes were bound by KAT7 in both cell lines (Fig. 4G). We then compared expression levels and KAT7 promoter occupancy of the spreading genes (Fig. 4H,I) and found that, consistent with the genome-wide patterns, OCI-AML3 showed good correlation between the two parameters (Fig. 4I). However, MOLM-13 did not show clear correlation, but KAT7 rather seemed to be highly bound to a small fraction of the MLL-AF9 spreading genes (Fig. 4H). We further compared KAT7 promoter occupancy with expression fold changes on day 3 after KAT7 KO and found that in MOLM-13 genes with higher KAT7 occupancy were more susceptible to and down-regulated upon KAT7 loss (Fig. 4J). In particular, *MEIS1, PBX3, JMJD1C* and *SENP6*, which were significantly down-regulated on day 3 (Fig. 3C), showed particularly high KAT7 promoter occupancy (Fig. 3J). There were significant differences in KAT7 promoter occupancy between the down-regulated and the up-regulated (n=4) or not-differentially expressed (n=126) genes (Fig. 3K). In sharp contrast and consistent with its KAT7 independence, in OCI-AML3 KAT7 loss did not show a significant impact on expression on day 3, even for genes with KAT7 promoter occupancy comparable to those down-regulated in MOLM-13 (Fig. 4L, M). In both cell lines, for non-MLL-AF9 target genes, there was no significant difference in KAT7 promoter occupancy between up-regulated, down-regulated and not-differentially expressed genes (Fig. S2A-D). Taken together, these results suggest that although KAT7 is not required for general transcription in AML, KAT7 and its acetylase activity are required for MLL-AF9 oncoprotein to sufficiently up-regulate and maintain expression of a specific set of the MLL-AF9 spreading genes.

**Figure 4.**
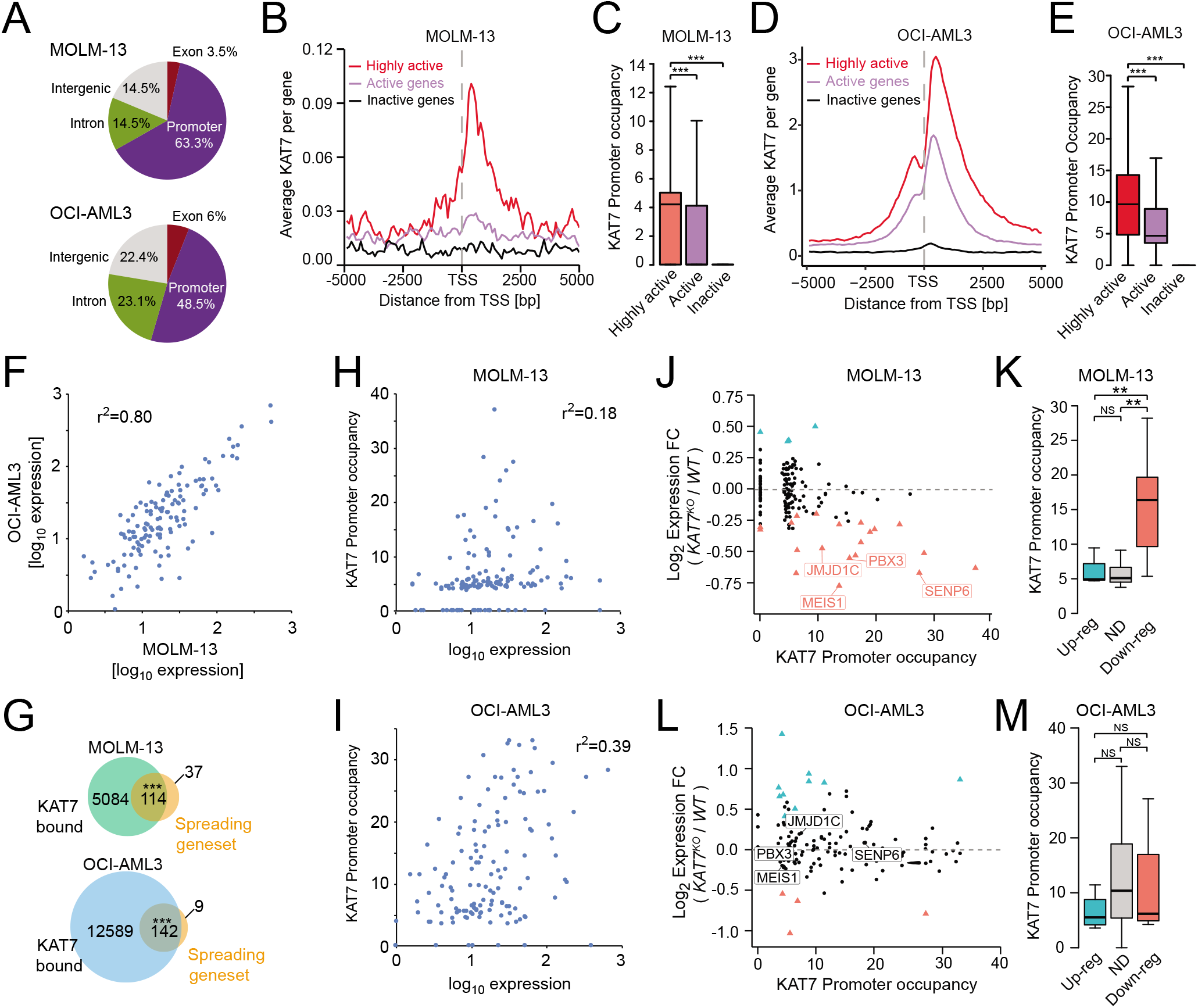
KAT7 binds to and is required for expression of a subset of MLL-AF9 targets. **A)** Distribution of KAT7 signals at promoters (TSS±2kb), exons (excluding promoter exons), introns and intergenic regions quantified by MACS2 peaks. **B)** Average occupancy of KAT7 per gene per base of highly active genes (>10 FPKM), active genes (>0, ≤10 FPKM) and inactive genes (FPKM=0) in MOLM-13. **C)** Box plots showing KAT7 promoter occupancy in highly active, active and inactive genes. Highly active genes showed significantly higher KAT7 promoter occupancy than active genes (*P* = 3.5 × 10^−227^) and inactive genes (*P* = 5.0 × 10^−324^) by Mann-Whitney-Wilcoxon test. **D-E)** Same as in B-C for OCI-AML3. Highly active genes showed significantly higher KAT7 promoter occupancy than active genes and inactive genes by Mann-Whitney-Wilcoxon test. **F)** Comparison of expression level of MLL-AF9 spreading genes between MOLM-13 and OCI-AML3. High correlation (r^2^=0.80) was observed. Expression data described in ref. 11 were used. **G)** Significant overlap between KAT7-bound and MLL-AF9-bound spreading genes in MOLM-13 (P = 3.9 × 10^−34^) and OCI-AML3 (P = 4.8 × 10^−15^) by Fisher’s Exact test. **H-I)** Comparison between expression and KAT7 promoter occupancy for MLL-AF9 spreading genes in MOLM-13 (**H**) and OCI-AML3 (**I**). Expression data described in ref. 11 were used. **J-M)** Comparison between KAT7 promoter occupancy and gene expression changes 3 days after KAT7 KO in MOLM-13 (**J-K**) and OCI-AML3 (**L-M**) for MLL-AF9 spreading genes. DE genes are highlighted in blue (up) or red (down). Down-regulated genes show significantly higher KAT7 promoter occupancy, compared to up-regulated (*P* = 0.0027) and not differentially expressed genes (*P* = 2.7 x 10^−7^) in MOLM-13 (**K**) but not in OCI-AML3 (**M**). One-tailed *ř*-test was performed.

### MLL-fusion associated machineries are evicted from their target loci upon KAT7 loss

To study the consequences of KAT7 loss at the chromatin level, we employed the auxin-inducible degron (AID) system ^39^. For this, we first stably expressed KAT7-AID and then knocked out the endogenous *KAT7* with gRNA. We confirmed that proliferation of KAT7-AID MOLM-13 was similar to that of parental MOLM-13 cells (Fig. S3A), indicating that AID-tagged KAT7 is functional. We then introduced OsTIR1 and treated cells with auxin indole-3-acetic acid (IAA) to deplete KAT7 protein. The KAT7 protein level rapidly decreased and plateaued 2 h after treatment (Fig. 5A). This was associated with complete loss of H3K14ac over the same time course (Fig. 5A) and followed by myeloid differentiation (CD11b) after 48 h (Fig. 5B). Therefore, the phenotype after IAA treatment faithfully mirrored those seen with CRISPR-mediated *KAT7* KO (Figs. 1C and 2A), indicating that the KAT7-AID system can be used to study the direct effects of KAT7 loss.

**Figure 5.**
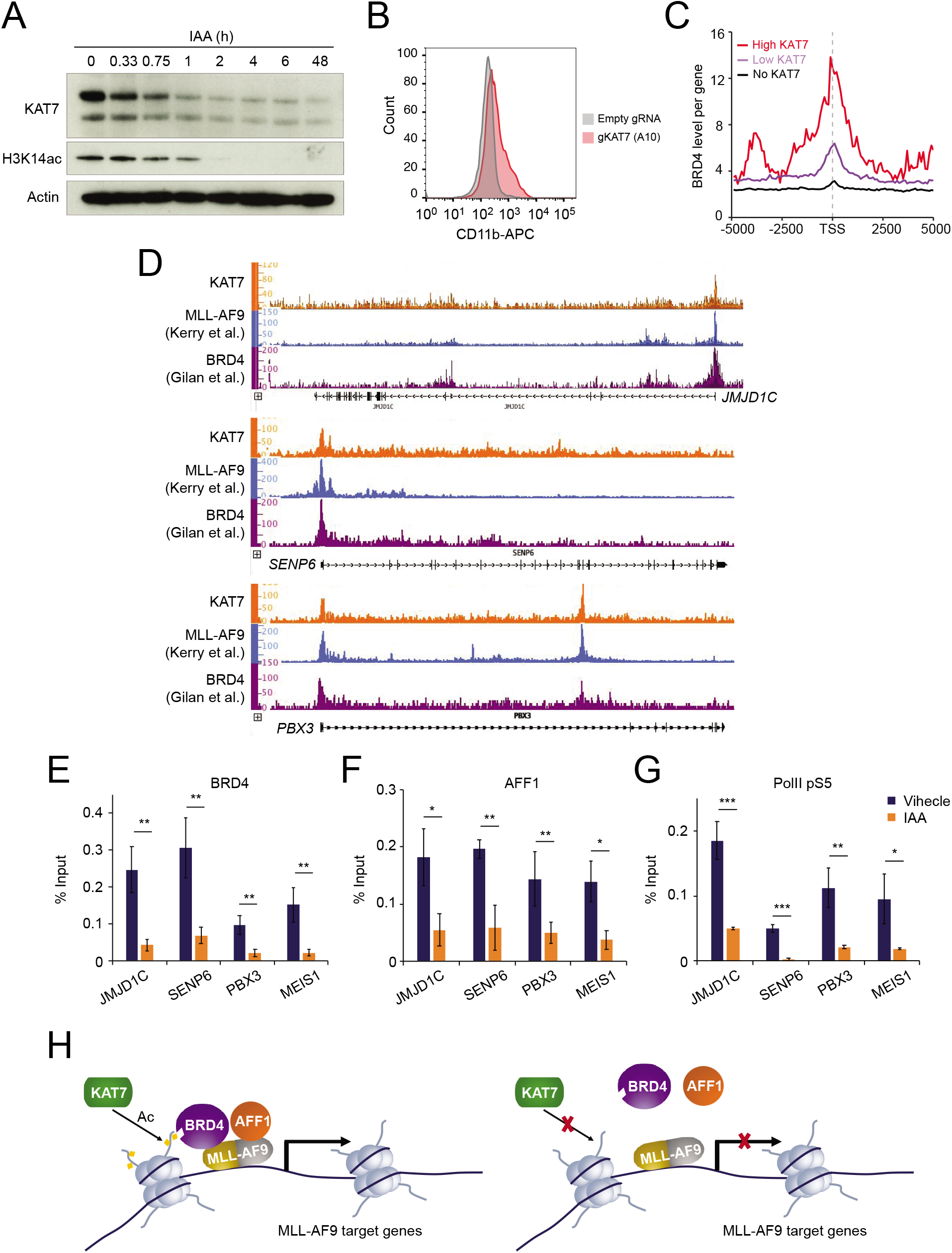
KAT7-dependent recruitment of BRD4 and SEC complex to a subset of MLL-AF9 spreading genes. **A)** Time-course western blot analyses of KAT7 and H3K14ac in IAA-treated KAT7-AID MOLM-13 cells. **B)** FACS analysis of IAA-treated KAT7-AID MOLM-13 at 24 h using CD11b antibody. **C)** Genome-wide correlation of KAT7 and BRD4 binding at promoter. **D)** Co-localization of KAT7, MLL-AF9 and BRD4 at the promoter of the key MLL-AF9 spreading genes. **E-G)** ChIP-qPCR analysis of BRD4, AFF1 and pS5 Pol II after 24 hours of IAA treatment at the promoter region of the key MLL-AF9 spreading genes. Data are shown as mean ± S.D. (n= 3). Two-tailed t-test was performed (*, *P ≤* 0.05; **, *P ≤* 0.01; ***, *P ≤* 0.001). **H)** Model of KAT7-dependent recruitment of BRD4 and SEC complex machinery to drive transcription of MLL-AF9 target genes.

Using this degron system, we first investigated whether KAT7 is required for MLL-AF9 fusion protein to maintain its binding to the promoters of key MLL-AF9 spreading genes, namely *JMJD1C, SENP6, PBX3* and *MEIS1*, which were down-regulated significantly on day 3 after *KAT7* KO (Fig. 3C) and bound by KAT7 with very high occupancy (Fig. 4J). ChIP followed by quantitative PCR (ChIP-qPCR) analysis using antibodies against MLL N-terminal and AF9 C-terminal regions 24 h after IAA treatment showed no significant changes in their occupancy at the promoter region (Fig. S3B). This observation is consistent with previous findings that MLL-AF9 recruitment is dependent on interactions with MENIN ^40^, LEDGF ^41^ and the PAF complex ^42^.

We next investigated the binding of the histone acetylation “reader” BRD4 at gene promoters. BRD4 is a key interactor of MLL-fusion proteins and a target of anti-leukemic drugs ^5^,^6^. Globally, BRD4 occupancy at promoter regions is highly correlated with KAT7 occupancy (Fig. 5C). At the selected key MLL-AF9 spreading genes, KAT7, MLL-AF9 and BRD4 highly co-localized (Fig. 5D). Upon KAT7 depletion, BRD4 markedly dissociated from the promoters of the key spreading genes (Fig. 5E). This was associated with dissociation of AFF1, a scaffold protein of the super elongation complex (SEC), from these promoters (Fig. 5F). In addition, RNA polymerase II Serine 5 phosphorylation (PolII-pS5) occupancy was also dramatically reduced at these loci (Fig. 5G). These results suggest that KAT7-mediated histone acetylation serves as a scaffold for BRD4 binding to chromatin, which is required for sustained recruitment of transcriptional activators (Fig. 5H).

## Discussion

Despite advances in understanding its genomics and molecular pathogenesis, AML remains lethal to the majority of sufferers ^43^ and anti-AML mainstream therapies have not changed significantly for several decades. Among different AML subtypes, cases driven by *MLL-X* fusion genes continue to represent a poor prognostic category ^1–3^ and despite recent developments in the field ^5^,^6^,^8^,^10^,^44^ clinical progress is still lacking, emphasizing the need for new therapies. Here, we demonstrate that KAT7 represents a promising novel therapeutic target for *MLL-X* AML and provide insights into its function in the maintenance of these leukemias.

KAT7-containing complexes have been shown to acetylates lysine residues of histone H3 and H4 tails. Specificity for the target histone is determined by the scaffold subunits, BRPF and JADE. In *in vitro* HAT assays using nucleosomes, BRPF-containing complex acetylates H3K14 and K23, while JADE-containing complex acetylates H4K5, K8 and K12 (ref. 23). These scaffold subunits can also form a HAT complex with KAT6A and KAT6B and show a similar specificity against H3 and H4 (ref. 23). In principle, this complex formation is interchangeable, but seems to be differentially regulated in different cell types. For example, in HeLa cells, purified KAT7-containing HAT complex contained JADE but not BRPF ^27^, and BRPF-containing HAT complexes predominantly contain KAT6A/B ^23^. In keeping with this, knock-down of *KAT7* in HeLa cells results in reduction of all of the three acetylation marks on H4 (ref. 27). In contrast, *KAT7*-KO in mouse embryonic fibroblasts and *KAT7* knock-down in erythroblasts led to reduction specifically of H3K14ac among the 5 potential acetylation sites and had no effect on histone H4 (ref. 24,25). In the present study, we found that KAT7 KO in AML cells resulted in complete loss of both H3K14ac and H4K12ac, a pattern that has not been described before. This strongly suggests that KAT7 is solely responsible for acetylation of these 2 sites and that KAT6A does not compensate for KAT7 loss. It has been shown that in mouse embryos KAT6A plays a specific role in regulating local H3K9 acetylation at the *Hoxa* and *Hoxb* loci and is not involved in global H3K9 and H3K14 acetylation ^45^. This might be the case in AML cells, indicating that KAT6A and KAT7 play non-overlapping roles in histone acetylation. KAT6A also showed fitness defects in *MLL-X* AML cell lines in our analysis. It would be interesting to investigate the molecular basis of KAT6A dependency and the mechanisms by which cell-type-specific complex formation is regulated.

Besides the general correlation between expression level and KAT7 promoter occupancy, we found that KAT7 strongly bound to a small subset of MLL-AF9 spreading genes in MOLM-13. The expression of this subset is especially susceptible to KAT7 loss and it includes well-known leukemia maintenance genes in *MLL-X*-driven AML such as *MEIS1, PBX3, JMJD1C* and *MEF2C*. MLL-X target genes are known to be associated with high level of H3K79me2, which is deposited by histone methyltransferase DOT1L^8,44^. In MLL-X spreading genes, H3K79me2 marks also spread into gene bodies, showing broader peaks ^38^. These spreading genes were shown to be more sensitive to DOT1L inhibition than the other genes (non-spreading and no-bound genes); the entire set of the spreading genes are downregulated upon DOT1L inhibition ^38^. This pattern of down-regulation is in sharp contrast to the case of KAT7 loss, in which only a subset of the spreading genes that are highly bound by KAT7 are down-regulated. Therefore, recruitment of KAT7 complex to promoters of this subset might be regulated differently from the rest of the promoters.

As a potential reader of H3K14ac and/or H4K12ac by KAT7, we investigated BRD4 and showed that BRD4 dissociated from the promoter of the key spreading genes upon KAT7 loss. BRD4 is known to play crucial roles in maintaining *MLL-X*-driven leukemic programs ^5^,^6^. A prominent role of BRD4 in AML is to activate super-enhancers at the *MYC* locus and maintain high *MYC* expression ^46^. Bromodomain inhibitors such as JQ1 and i-BET151 disrupt the interaction between BRD4 and acetylated histones, and evict BRD4 from chromatin, thereby leading to rapid downregulation of *MYC*. In addition to the role at enhancers, BRD4 is known to play a role in promoting Pol II elongation at promoter-proximal regions by recruiting SEC including P-TEFb ^46^. Accordingly, we observed the reduced binding of AFF1 scaffold protein of SEC upon KAT7 loss. This suggests that Pol II elongation is severely affected, causing downregulation of target genes. However, since the role of BRD4 at promoters is generic, it is still unclear why only a small subset of MLL-AF9 spreading genes are affected by KAT7 loss in MOLM-13 and why gene expression is relatively stable, even in the absence of KAT7, in the KAT7-independent OCI-AML3 cells. It is plausible that additional acetylated histone readers are involved in KAT7-dependent transcriptional activation for the MLL-AF9 spreading genes. From this perspective, it is interesting to observe that pS5-Pol II reduced its occupancy at the selected promoters. Serine 5 of Pol II is phosphorylated by TFIIH after Pol II loading onto promoters ^47^. Subsequent Serine 2 phosphorylation on the C-terminal domain of Pol II by P-TEFb promotes Pol II elongation ^47^. Therefore, KAT7-mediated histone acetylation may also be required for the initiation phase of Pol II transcription. Further studies are required to elucidate the molecular basis of KAT7-dependent transcription of MLL-AF9 target genes.

Collectively, our findings reveal that KAT7 acts upstream of BRD4 to recruit MLL-fusion associated machineries to the promoter of key MLL-AF9 target genes, which represent an alternative therapeutic target for *MLL-X* leukemia. We anticipate that our work will motivate the development of small-molecule inhibitors that are specific to KAT7 and highlight KAT7 as a potential therapeutic target for MLL-fusion AML.

## Acknowledgements

This work was funded by the Wellcome Trust (WT206194), the Kay Kendall Leukaemia Fund (KKL920), Bloodwise (17006) and Exonate Ltd. K.T. was funded by a Wellcome Trust Sir Henry Wellcome Fellowship (grant reference RG94424). G.S.V. was funded by a Cancer Research UK Senior Cancer Fellowship (C22324/A23015) and a Wellcome Trust Senior Fellowship in Clinical Science (WT095663MA). We thank Bee Ling Ng, Jennifer Graham, Christopher Hall and Sam Thompson from the Wellcome Sanger Institute Cytometry Core Facility team for help with flow cytometry. We are grateful to the staff of the Sanger Institute Research Support Facility for help with mouse experiments and the staff of the Sanger Institute Core Sequencing facility for sequencing. We thank Mathew Garnett for providing the Nomo-1 cell line; Pedro for his help compiling figures using Adobe Illustrate and Josep Nomdedeu for help advices in writing the manuscript.

## Authors contributions

K.Y. conceived the study and designed the experiments. Y.Z.A performed the experiments and analyzed the data. E.D.B. performed and analyzed the in vivo mouse studies. M.G. and S.H.O. conducted bioinformatic analyses. J.Y., B.J.P.H., M.G., and K.T. helped with data interpretation and direction. M. Gozdecka and X.C. advised on ChIP experiments. Y.Z.A., M.G., J.Y., G.S.V. and K.Y. wrote the manuscript. All authors reviewed the manuscript.

